# Ultrafast Brownian-ratchet mechanism for protein translocation by a AAA+ machine

**DOI:** 10.1101/2020.11.19.384313

**Authors:** Hisham Mazal, Marija Iljina, Inbal Riven, Gilad Haran

## Abstract

AAA+ ring-shaped machines, such as ClpB and Hsp104, mediate substrate translocation through their central channel by a set of pore loops. Recent structural studies suggested a universal hand-over-hand translocation mechanism, in which pore loops are moving rigidly in tandem with their corresponding subunits. However, functional and biophysical studies are in discord with this model. Here, we directly measure the real-time dynamics of the pore loops of ClpB and their response to substrate binding, using single-molecule FRET spectroscopy. All pore loops undergo large-amplitude fluctuations on the microsecond timescale, and change their conformation upon interaction with substrate proteins. Pore-loop conformational dynamics are modulated by nucleotides and strongly correlate with disaggregation activity. The differential behavior of the pore loops along the axial channel points to a fast Brownian-ratchet translocation mechanism, which likely acts in parallel to the much slower hand-over-hand process.

---

AAA+ (ATPases associated with various activities) proteins form an abundant family of biological machines that harness the energy of ATP binding and hydrolysis to power cellular tasks such as protein unfolding, protein disaggregation, DNA helicase activity, DNA replication initiation, and cellular cargo transport (*1, 2*). These machines typically assemble into hexameric ring complexes that envelope a large central channel (*3*). Pore loops lining the axial channel are essential elements for machine activity (*3, 4*), and may exert force to pull substrates through the channel. The remarkable disaggregation machine, ClpB, and its eukaryotic analog Hsp104 consist of two nucleotide binding domains (NBDs) per subunit. NBD1 contains two functional pore loops, PL1 and PL2, while NBD2 contains a single functional pore loop, PL3 (*5–8*) (Fig. 1). Both PL1 and PL3 harbor a conserved motif that involves a functionally important tyrosine residue, and their role in substrate pulling in a nucleotide-dependent manner has been shown (*6, 7, 9–11*).

**Fig. 1.**
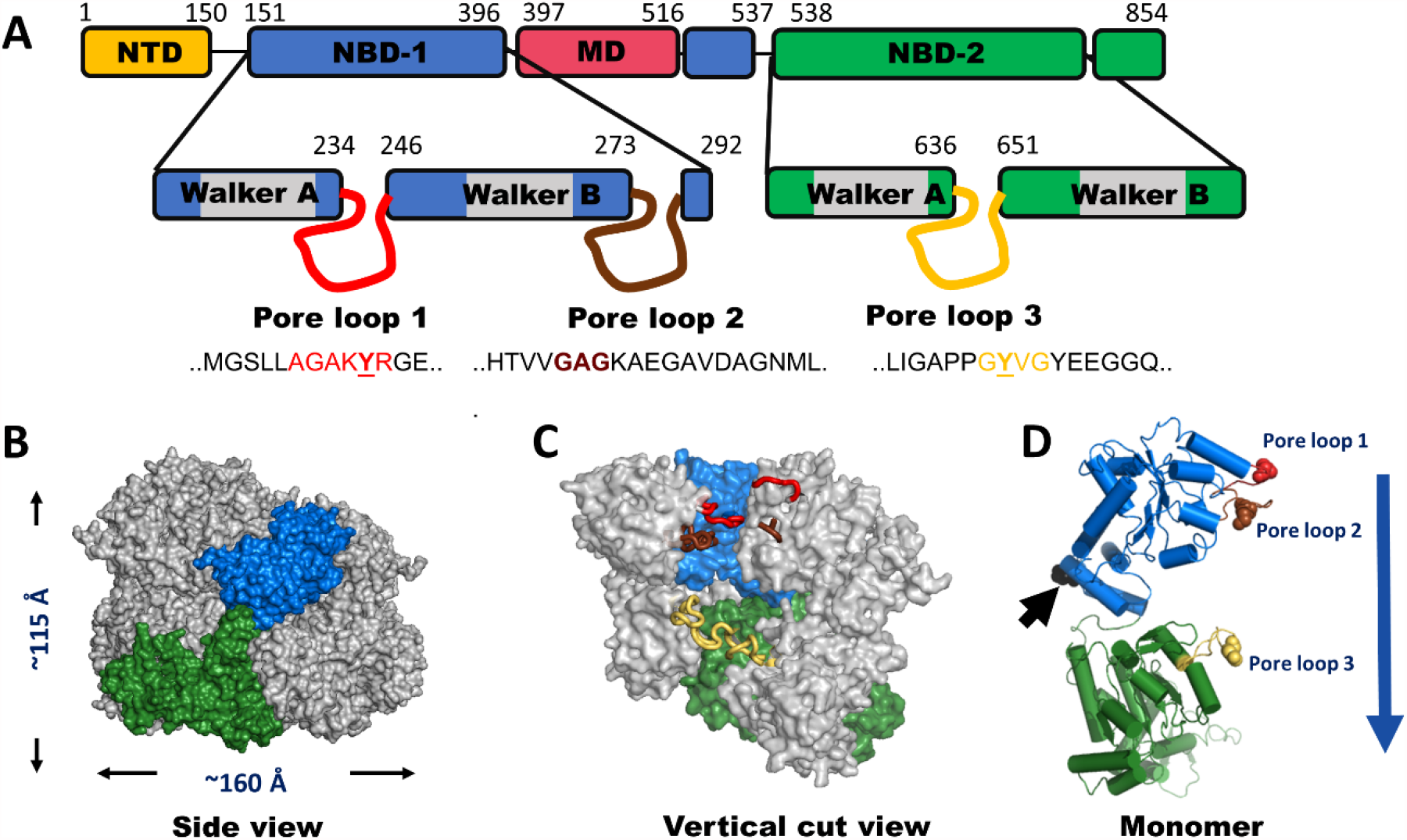
ClpB domain organization and structure. (A) Domain organization of the monomer of ClpB. NTD- N-terminal domain. NBD- nucleotide binding domain. MD- middle domain. The location and sequence of pore loops are indicated on each NBD. The numbers mark residue positions in the sequence. A ClpB variant that lacks the NTD is used in this study. This variant actually occurs naturally and is fully functional. (B) Side view of the hexameric model of *E. coli* ClpB reconstructed from cryo-EM data (PDB: 6OAX). One protomer (protomer A) is highlighted using the same color code as in A. The MD and the NTD are not resolved in this structure, thus they are not shown. (C) Vertical cut view of the hexameric structure in (B). PL1, PL2 and PL3 of one protomer are colored in red, brown and yellow, respectively. PL1 and PL2 are not fully resolved. (D) The monomeric structure of *E. coli* ClpB (PDB: 6OAX, protomer A), *α*-helices are represented as cylinders, pore loops are shown in the same color as in C. The spheres on pore loops represent the conserved tyrosine residues of PL1 (red) and PL3 (gold), and the alanine residue of the motif GAG in PL2 (brown). Black arrow points to the position P368, which is equivalent to position S359 in *TT* ClpB, used in our smFRET experiments. Blue arrow indicates the proposed direction of substrate translocation.

Recently, high-resolution structures of multiple AAA+ molecular machines were solved by cryogenic electron microscopy (cryo-EM) (*3, 12–15*), showing that these proteins form hexameric spirals (Fig. 1 B-C), rather than the fully-symmetric hexameric rings described earlier (*16*). Based on these recurrent features, it has been proposed that substrate-polypeptide threading through the central channel of the machines proceeds in a sequential, hand-over-hand manner, facilitated by rigid-body movement of the protomers with a step size equivalent to two amino acids (aa) per ATP hydrolysis cycle (*17, 18*). Since the ATP hydrolysis rate of these proteins is low, on the order of ~ 0.05 – 3.5 ATP molecules per second (*12, 13, 15, 19*), the translocation rate is expected to be slow. However, a recent single-molecule force spectroscopy study showed that, remarkably, translocation by ClpB is extremely fast, ~ 240 – 450 aa per second, and occurs in bursts of 14 – 28 aa (*20*). Studies on ClpX (*21*) and ClpA (*22*) also demonstrated substrate translocation with step sizes larger than 2 aa. Furthermore, it was found that these machines were active in translocation even when several of their six subunits were rendered inactive (*23, 24*). The strong disagreement between models based on structural studies and the results of real-time measurements is intriguing and hinders our understanding of the basis of substrate remodeling by these machines.

A molecular picture of the elusive pore-loops motions is critical for elucidating the translocation mechanism of such complex machines. In this work, we tackle this problem by directly observing the dynamics of pore loops in ClpB and their coupling along the axial channel, as well as their direct coupling during substrate translocation, using single-molecule FRET (smFRET) experiments (*25, 26*). First, we studied PL1, which is located on NBD1 (Fig. 1 D). To probe PL1 dynamics in its complete functional form, we mutated and labeled several residues flanking its conserved motif “AGAKYR” (Fig. 1 A). We found that only the variant S236C maintained disaggregation activity after labeling (Fig. 2 A, Fig. S1-3, Table S1). We picked residue S359, located at the center of the vertical axis of the ClpB protomer, as a reference point to probe pore-loop motions along the axial channel (Fig. 1 D). The same reference point was also used to study motion of the other pore loops. We thus prepared and labeled the variant S236C-S359C with donor (Alexa 488) and acceptor (Alexa 594) dyes, and assembled ClpB molecules such that only a single subunit within each ClpB hexamer was labeled. Fluorescence anisotropy measurements (Fig. S4A, Table S2) ruled out motional restrictions of the fluorescent dye induced by interaction with ClpB’s channel or with substrate proteins (see below).

**Fig. 2.**
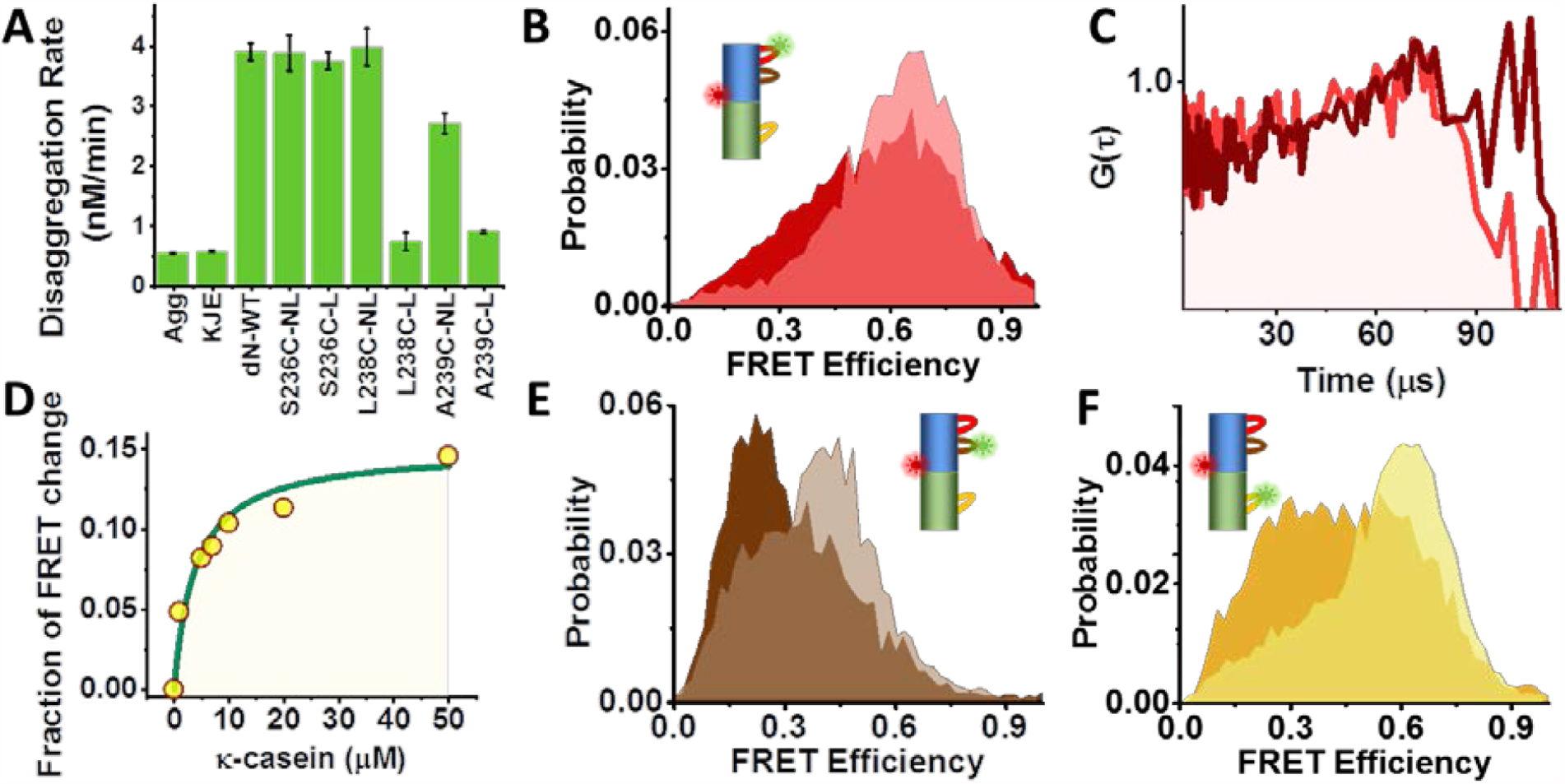
Probing pore-loop conformational dynamics using smFRET: (A) Disaggregation activity of ClpB WT and PL1 mutants. Results show that all non-labeled pore loop mutants (NL) were active, but only the mutant S236C maintained activity after labeling (L). (B) FRET efficiency histograms of the PL1 variant without substrate (light red), and in the presence of 20 µM κ-casein (dark red). (C) Donor-acceptor fluorescence cross-correlation functions of the PL1 variant; light red- without substrate, dark red- with substrate. (D) The relative change of the low FRET efficiency population in histograms (Fig. S6) as a function of κ-casein concentration. (E) FRET efficiency histograms of the PL2 variant without substrate (light brown) and in the presence of 20 µM κ-casein (dark brown). (F) FRET efficiency histograms of the PL3 variant without substrate (light orange) and in the presence of 20 µM κ-casein (dark orange).

Photon-by-photon FRET trajectories were collected from labeled ClpB molecules freely diffusing through a focused laser beam in the presence of 2 mM ATP. The FRET efficiency histogram constructed from the data (Fig. 2 B, Fig. S5 A) showed a major population at a FRET efficiency value of 0.65 ± 0.01. This major peak was found to be much broader than expected based on shot noise, indicating dynamic heterogeneity (see also Fig. S5 B). Donor-acceptor fluorescence cross-correlation calculated from the same data showed a rising component on a time scale of tens of microseconds, symptomatic of fast dynamics (Fig. 2 C). To understand the role of PL1 in substrate engagement, we studied the effect of the soluble model substrate κ-casein (*19, 27*). Enhanced ATPase activity of ClpB was registered in the presence of the substrate (Fig. S2 B), and its translocation was verified using BAP-ClpP system (Fig. S1C). Importantly, in the presence of 20 µM κ-casein we noted a substrate-concentration dependent shift of the FRET efficiency histograms toward lower values (Fig. 2 B, Fig. 2 D, Fig. S6), indicating a major conformational change, though no significant substrate-induced change in the dynamics was seen in the cross-correlation function (Fig 2 C).

To study PL2 dynamics in a similar manner (Fig. 1 D), we mutated an alanine residue at position 281 to cysteine. The measured FRET efficiency histogram of the double-labeled A281C-S359C mutant was dramatically changed upon the addition of 20 µM κ-casein (Fig. 2 E). The donor-acceptor fluorescence cross-correlation function again pointed to microsecond dynamics (Fig. S7 A). To study PL3, we mutated Y646 close to the conserved ‘GYVG’ motif. The FRET efficiency histogram of labeled S359C-Y646C in the presence of 2 mM ATP (Fig. 2 F, Fig. S5 E) changed significantly upon the addition of 20 µM κ-casein, with a substantial increase in the low FRET efficiency shoulder. Fluorescence cross-correlation analysis showed again evidence for fast dynamics (Fig. S7 B). We ruled out the involvement of NBD1-NBD2 interdomain motion in these dynamics (Fig. S8).

To gain further insight into pore-loop dynamics, we used H^2^MM, a hidden Markov model algorithm for photon-by-photon analysis capable of identifying microsecond motions (*28*). A continuous representation of the free-energy profile of each pore loop was obtained by using a model of 9-10 equally spaced and sequentially connected states. Interestingly, two well-defined potential wells, rather than a single minimum, were retrieved in each case (Fig. S9), with microsecond time-scale jumps between them. We therefore continued to model the data with two states, globally analyzing pairs of data sets measured with and without κ-casein. From this analysis, validated using several methods (Figs. S10-12, Tables S3-S5), we obtained the population of each pore loop in its two states with and without substrate. An effective equilibrium coefficient for the conformational dynamics of pore loop *i*, 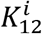, was defined as its population ratio. Thus, for example, the population ratio in pore loop 1, 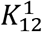 was 0.48 ± 0.01, and changed to 0.81 ± 0.01 in the presence of κ-casein (Fig. 3 A, see Table S6 for all 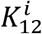 values). The microsecond rates of transition between states obtained from the analysis were in good agreement with the cross-correlation results (Table S3). We utilized state FRET-efficiency values obtained from the H^2^MM analysis, together with information from ClpB structures (Table S4, Fig. S13 A) to estimate the amplitude of motion of each pore loop. We calculated a motion of more than 10 Å in all cases, corresponding to as much as two substrate-protein residues. These large fluctuations, which could not be inferred from recent static high-resolution structures (Fig. S13 B-D), are likely to contribute to substrate translocation on a much faster timescale than expected based on ATP hydrolysis rates.

**Figure 3.**
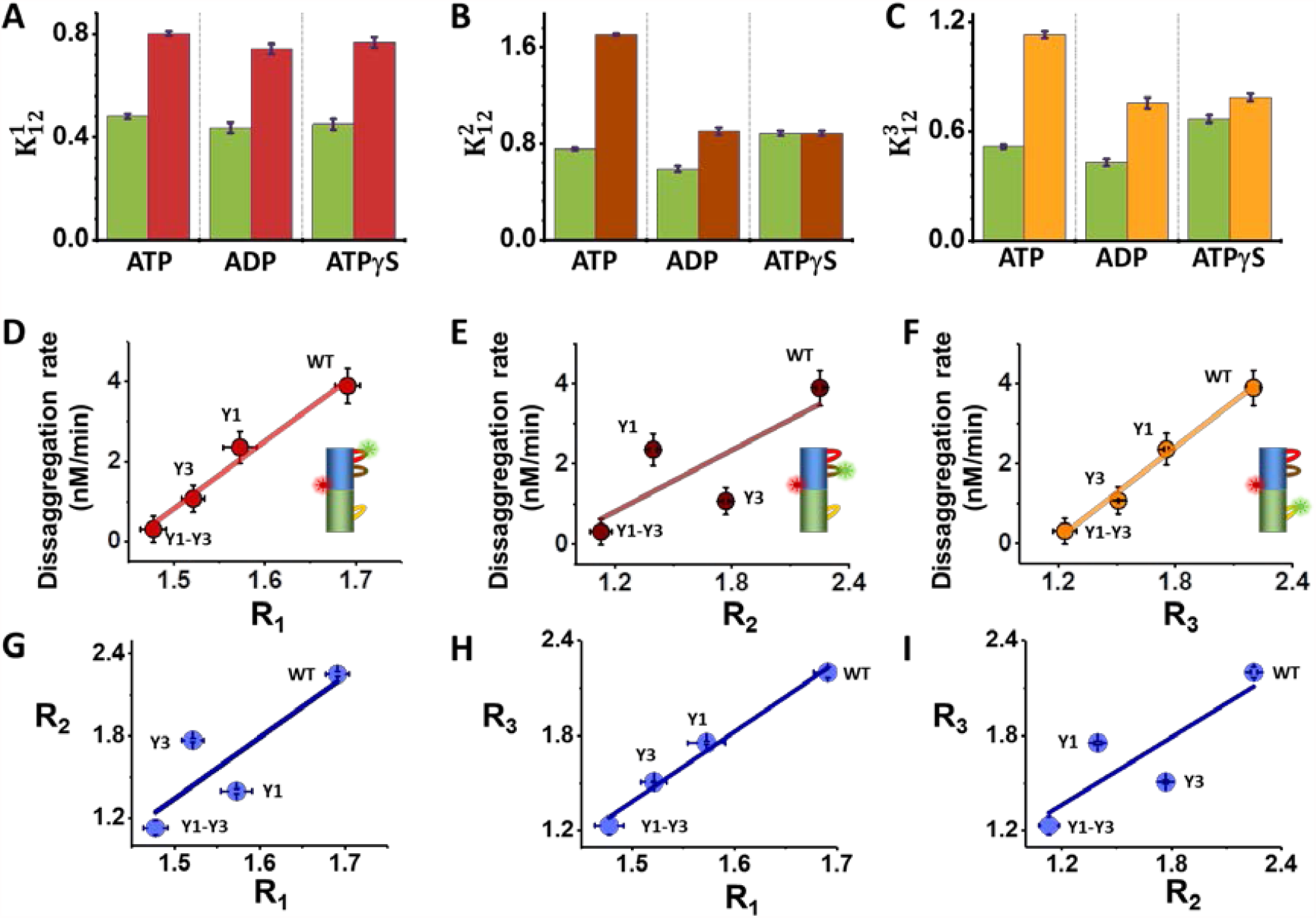
Modulating pore loop conformational changes by nucleotides and tyrosine mutations. (A-C) Equilibrium coefficients for the conformational dynamics without or with 20 µM κ-casein and in the presence of different nucelotides. (A) PL1: green- no substrate, red- with substrate. (B) PL2: green- no substrate, brown- with substrate. (C) PL3- green- no substrate, yellow- with substrate. (D-F) Correlation of disaggregation rate and the substrate-response factors, *R*_*i*_, of the three pore loops in tyrosine mutants. (G-I) Correlations between substrate-response factors of the three pore loops in tyrosine mutants. Errors were calculated from at least two experiments.

To understand the mechanochemical coupling between ATP hydrolysis and pore-loop motions, we studied the effect of different nucleotides on the dynamics, including (in addition to ATP) ADP to mimic the post-hydrolysis state and the slowly hydrolysable analog ATPγS. To quantify the effect of nucleotides on pore-loop conformational dynamics, we computed 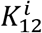 values from the H^2^MM results (Table S6) and then calculated the substrate-response factors, which are ratios of 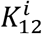 values with and without the substrate, 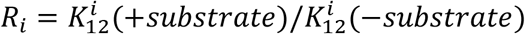 (Table S8). Surprisingly, FRET efficiency distributions of PL1 registered in the presence of ADP, and ATPγS, were almost identical to the equivalent histograms measured with ATP, either with or without κ-casein (Fig. S14 A-B). Correspondingly, 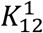 values (Fig. 3 A) were very similar irrespective of the nucleotide used, and so were *R*_*1*_ values (Table S8): 1.68 ± 0.01 with ATP, and 1.69 ± 0.02 with ADP and ATPγS. In contrast, PL2 demonstrated a differential response to nucleotides (histograms in Fig. S15 C-D, 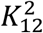 in Fig. 3 B). Accordingly, *R*_*2*_ changed from 2.25 ± 0.01 in ATP to 1.53 ± 0.02 in ADP, indicating a weaker response to the substrate. Remarkably, ATPγS did not elicit any substrate-induced change in the FRET efficiency histogram of PL2, and *R*_*2*_ for this nucleotide was 1.00 ± 0.02.

In the case of PL3, the FRET efficiency histograms (Fig. S14 E-F) and 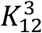 values (Fig. 3 C) also showed the largest response to substrate addition in the presence of ATP, with a corresponding *R*_*3*_ value of 2.20 ± 0.02, though significant changes were also registered in the presence of ADP (*R*_*3*_ = 1.76 ± 0.02), and almost no response with ATPγS (*R*_*3*_ = 1.17 ± 0.02). Thus, we can conclude that ATP hydrolysis and likely the presence of the product P_i_ are required for a significant shift of the free energy surface of PL2 and PL3 by the substrate, while PL1 is completely nucleotide-type independent.

To probe the correlation of pore-loop conformational changes with machine activity, we investigated mutants of the conserved tyrosine residues located on PL1 and PL3. Several studies showed that mutation of these tyrosine residues to alanines resulted in reduced machine activity, with a more pronounced effect in the case of PL3 (*6, 11, 29*). We generated either a tyrosine mutant of PL1, Y243A (Y1), a tyrosine mutant of PL3, Y643A (Y3), or a double-mutant, Y243A-Y643A (Y1-Y3). G6PDH disaggregation activity was reduced in all tyrosine mutants (Fig. S2 C, Table S9) (*11, 29*), with Y3 showing a stronger activity reduction than Y1, and Y1-Y3 showing almost no disaggregation activity. smFRET measurements were performed on each pore loop in constructs bearing Y1, Y3 or Y1-Y3, in the presence of 2 mM ATP and with or without 20 µM κ-casein (Figs. S15-16, and Tables S7-8). Correlation analysis of the disaggregation activity of each tyrosine mutant with substrate-response factors, *R*_*i*_ (Fig. 3 D-F), demonstrated significant correlation for PL1 and PL3, with *R*^*2*^ values of 0.98 and 0.95, respectively. In contrast, PL2 demonstrated a weaker correlation, with an *R*^*2*^ value of 0.64. These results indicate that the conformational changes occurring in both PL1 and PL3 contribute significantly to machine activity. The high correlation between PL3 dynamics and the overall disaggregation activity of the machine is in line with previous findings that NBD2 is the main contributor to machine activity (*19, 30*), but the high correlation of PL1 dynamics to disaggregation is less anticipated. Strikingly, correlation plots between substrate-response factors of the different pore loops (Fig. 3 G-I) demonstrated a strong correlation between PL1 and PL3, with an *R*^*2*^ value of 0.99. All correlations observed here were validated through a calculation that did not depend on H^2^MM analysis of the data, as described in the Methods section.

Our findings confer a more active role to pore-loop dynamics in machine activity than perceived before. Indeed, the microsecond conformational dynamics of PL1 and PL3 are not only correlated with each other, but also strongly correlate with the disaggregation rate of the machine, and therefore also with substrate translocation rate. At the same time, both PL2 and PL3, but not PL1, require nucleotide hydrolysis to change their conformation in response to substrate translocation, even though their dynamics are much faster than ATP hydrolysis.

These remarkable results point to a Brownian-ratchet like mechanism (*31–33*) for fast substrate translocation by ClpB. In a Brownian ratchet (Fig. 4 A), the input of chemical energy (e.g. ATP hydrolysis) transfers the machine between a state that allows free diffusion and a state with a pawl-like free-energy surface that rectifies the overall motion. The strong correlation between the structural changes of PL1 and PL3 and the disaggregation rate implies that these pore loops are active in pulling the substrate across ClpB’s channel (Fig. 4 B). At the same time, the requirement for ATP hydrolysis for PL2 and PL3 conformational changes points to these pore loops as the ratchet pawls that operate to rectify substrate motion at different stages of translocation through the channel. PL2 acts first as a pawl when the substrate interacts with NBD1 and ATP has been hydrolyzed (Fig. 4B step 3). Similar events at NBD2 then engage PL3 as a pawl (step 4). Disengagement of these pawls may allow looped polypeptide segments to escape after partial threading (*20, 34*). The protein harnesses the power of asynchronous pulling by neighboring subunits to generate rapid processive translocation events. The fast fluctuations of the pore loops allow them to reconfigure along a protein substrate, facilitating proper gripping and pulling and likely preventing premature stalling. This Brownian ratchet can operate in parallel to the much slower hand-over-hand process, and is likely a general mechanism in AAA+ proteins.

**Figure 4.**
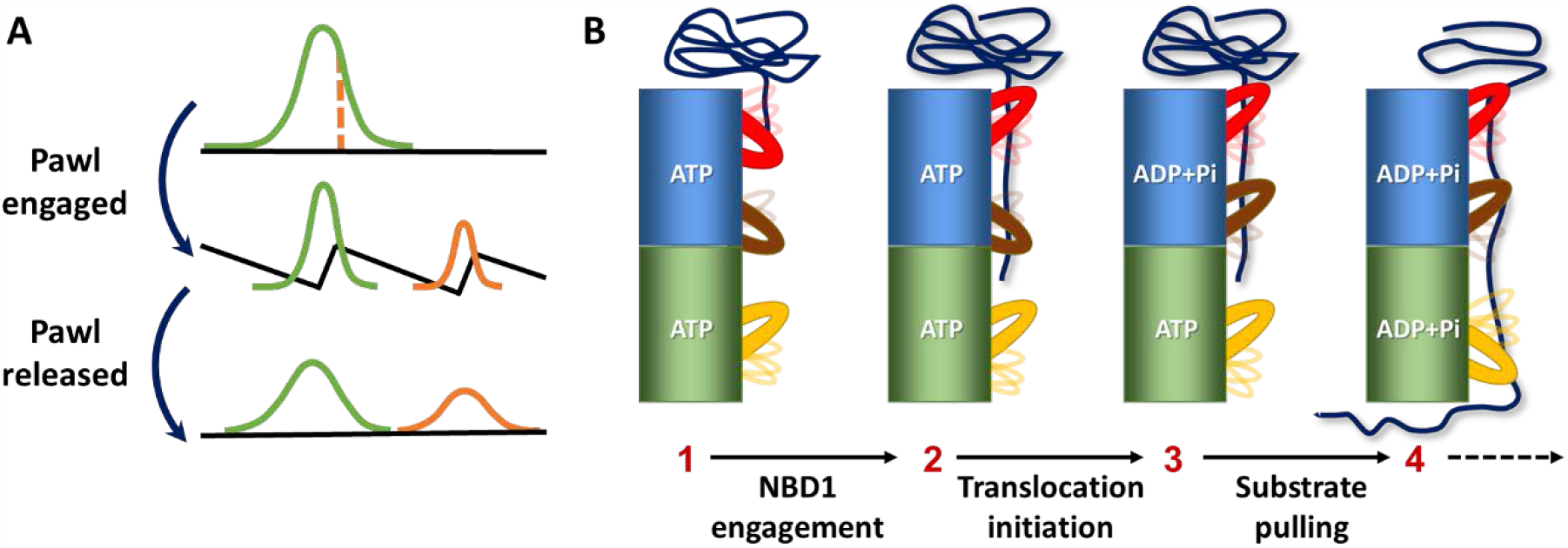
Brownian-ratchet mechanism for fast substrate translocation by ClpB. **A**. In a Brownian ratchet, an effective pawl periodically switches molecular dynamics between a flat free-energy surface and a structured surface, promoting unidirectional motion. **B**. Model for the Brownian-ratchet action of pore loops. As a substrate is engaged, pore loops gradually change their average conformation even while continuing to fluctuate on the microsecond time scale. Conformational changes in PL2 and PL3 take place only upon hydrolysis of ATP, turning them into effective pawls. At the same time, PL1 and PL3 function in pulling the substrate through the central channel.

## Acknowledgements

We thank Dr. Rina Rosenzweig for the generous gift of the BAP-ClpP plasmids and for helpful discussions. We further thank Drs. Hagen Hofmann and Pierre Goloubinoff for their careful reading of the manuscript.

## Author contributions

H.M. and G.H. conceived the project. H.M and and M.I. performed the experiments with help from I.R. All authors wrote the manuscript.

## Data and materials availability

the authors will make any data set and material generated in this work available to any researcher upon a reasonable request.

## Supplementary Materials

Methods. Figs. S1-S17.

Tables S1-S9.

